# Visual Illusion: A Matter Of Disorder Between Retinal And Perceived images

**DOI:** 10.64898/2026.11.14.766692

**Authors:** Avijit Paul, Amrita Mukherjee

**Affiliations:** Department of BME, Tufts University, USA; CSCR, ISI Kolkata, India; Applied Sciences, IIIT Allahabad, India

## Abstract

Visual illusions reveal that perception is constructive rather than a faithful copy of the retinal image. This paper proposes that the brain supplements ambiguous or incomplete sensory input with virtual edges, filled-in surfaces, brightness and size cues, and illusory contours. Using gray-level Shannon entropy, we compared original and perceived images across five illusion classes: edge detection, surface filling-in, brightness perception, size perception, and the Hermann grid. In all cases, perceived-image entropy exceeded real-image entropy. These findings support a unified account of visual illusions as constructive inference: the brain introduces additional information to achieve global coherence, reflecting a trade-off between low-entropy processing and perceptual interpretation.

## INTRODUCTION

The most important thing about human vision is that visual perception is similar to remote sensing or exteroceptive perception. Our eyes, along with the brain, help us to perceive the surrounding environment visually. At the back of our eyes, there is a layer of photoreceptors, rods and cones, known as the retina. Light from real images falls on the retina (equipped with 260 million photoreceptors), and the signals produced by those images pass through only 2 million ganglion cells before exiting via the optic nerve and reaching the primary visual cortex of the brain. With some visual processing in the visual cortex, from V1 and V2 to higher visual areas like the inferotemporal cortex [1, 2], the images we perceive can be thought of as newly reconstructed, cloned images that are pictorially similar to the originals. There are some apparent differences between real and perceived images that can be termed as visual illusions. Human beings have been fascinated with various kinds of optical illusions since ancient times. Several illusions, such as brightness contrast, brightness assimilation, amodal completion, size perception, neon-color spreading, Herman grid, etc., draw the attention of psychologists and neuroscientists to infer the logic of those visual images. Some scientists tried to describe particular illusions using their preferred logic [3]. Despite the reasonableness of those explanations, there is no generic logical deduction for different illusions to date. We derived a common logic underlying the visual illusions in this paper.

## IMAGE ENTROPY

Entropy is a physical state function that was first described by Rudolf Clausius [4] for isolated systems in thermodynamic equilibrium, in relation to the 2nd law of Thermodynamics. In the 1870s, Ludwig Boltzmann developed the concept of entropy from the statistical behavior perspective [5], which can be visualized from Equation 1.

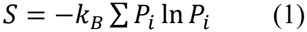

Where, *k*_*B*_ is the Boltzmann constant, and *P*_*i*_ is the probability of the system being in ith state. Many years later, in 1948, Claude Shannon conceptualized information entropy as the measure of disorder or uncertainty [6]. Shannon’s entropy is shown in Equation 2.

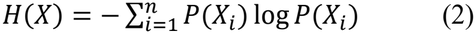

In a general sense, entropy is a measure of disorder in the whole system. For example, an ordered crystal has low entropy while its entropy increases as it is heated into a liquid state. Entropy is directly proportional to the number of states. In an image matrix, the grey levels correspond to different states, so an 8-bit pixel system has 28 states. In a nutshell, the entropy (H) of an image [7] is defined as:

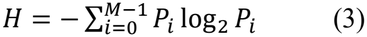

Where, *M* is the number of grey levels, and *P*_*i*_ is the respective histogram count probability associated with level *i*.

## PROPOSED WORK AND RESULTS: VISUAL ILLUSIONS AND IMAGE

### 3.1 ENTROPY

Our body always wants to be in a low-entropy state because more ordered structures are less vulnerable to decay [8]. Likewise, our brain always tries to process low-entropy signals. We human beings not only gather local information from an image but also try to churn out global patterns from some images to perceive their significance. This also differentiates us from chimpanzees, who are very good at quickly finding local features. To get the essence of images, our brain compares current images with preloaded data / shapes in memory. Then, in some of the cases, the brain inserts certain disorder as a form of extra information along with the original retinal input. In this way, the perceived images’ entropy increases. So whatever illusions occur in visual phenomena, all have similar characteristics: To perceive the retinal images, the brain inserts some useful extra information to create higher entropy images that are more acceptable to us. To calculate the entropy of an image, we used entropy() function of MATLAB R2010a (Version: 7.10.0.499). In the following sections, we will explore the analysis of entropy for visual illusions from different perspectives, such as:

- Edge Detection
- Surface Filling-in
- Brightness Perception
- Size Perception
- Hermann Grid Illusion.

The relevant figures present bar diagrams comparing the entropy of the images with that of the perceived illusions. Tables 1 through 5 list the entropy values for the various images and their corresponding perceived illusions, computed using MATLAB *entropy*() function. These tables clearly show that the perceived entropy exceeds that of the actual retinal image.

**Table 1:** Illusory Edge Detection.

| Visual Illusion | Real image Entropy | Perceived image Entropy |
|---|---|---|
| Kanizsa Triangle | 0.242846263527603 | 0.302962066055277 |
| Spoke-Circle Illusion | 0.355416571247298 | 0.395696580814625 |
| Amodal Completion | 1.889878630137753 | 1.941830818086904 |

**Table 2:** Illusory Surface Filling-in.

| Visual Illusion | Real image Entropy | Perceived image Entropy |
|---|---|---|
| Watercolor Illusion | 1.935959033222966 | 3.095444623219728 |
| Neon Color spreading | 1.161482965945826 | 1.651568130278114 |

**Table 3:**
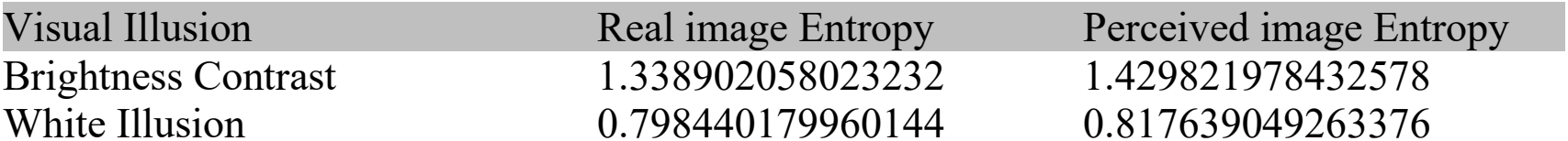
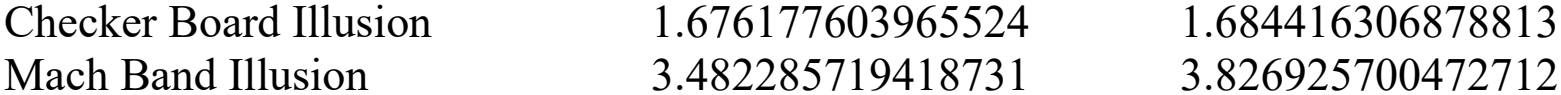
Brightness Perception.

| Visual Illusion | Real image Entropy | Perceived image Entropy |
|---|---|---|
| Brightness Contrast | 1.338902058023232 | 1.429821978432578 |
| White Illusion | 0.798440179960144 | 0.817639049263376 |
| Checker Board Illusion | 1.676177603965524 | 1.684416306878813 |
| Mach Band Illusion | 3.482285719418731 | 3.826925700472712 |

**Table 4:** Illusory Size Perception.

| Visual Illusion | Real image Entropy | Perceived image Entropy |
|---|---|---|
| Vertical-Horizontal Illusion | 0.126412568258541 | 0.136910972506488 |
| Curved Bar Illusion | 0.227486422674187 | 0.249293356331464 |

**Table 5:**
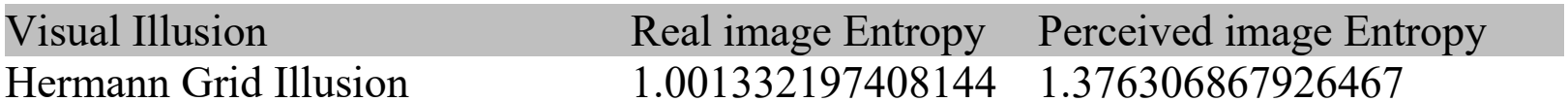
Hermann Grid Illusion.

| Visual Illusion | Real image Entropy | Perceived image Entropy |
|---|---|---|
| Hermann Grid Illusion | 1.0013321974081­44 | 1.376306867926467 |

### 3.2 EDGE DETECTION

To segregate the foreground image from the background where brightness differs significantly from the neighbors, while doing edge detection, our brain demarcates an outer boundary for every object to perceive it separately from the background. While real retinal inputs may lack edge information, or images explicitly do not display edges, our brain still adds virtual edge information, which ultimately increases the entropy of the perceived images. Some of the illusory edge detection phenomena include the Kanizsa Triangle [9], the Spoke-Circle illusion, and Amodal completion [10], etc. For all of these incidents, our brain probably takes the locally separated information into consideration to find a comprehensible pattern that is logically derivable from past experiences. To create this kind of virtual reality, the perceived image entropy increases. The following figures (Figure 1 (Left): Kanizsa Triangle illusion and Figure 1 (Right): Spoke-Circle illusion and Figure 2: Amodal completion) show the entropy measurements.

**Figure 1:**
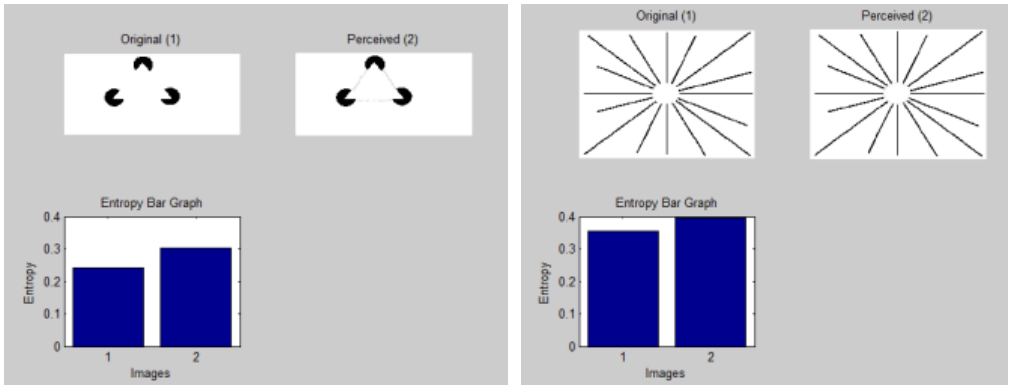
Kanizsa Triangle illusion (Left) & Spoke-Circle illusion (Right)

**Figure 2:**
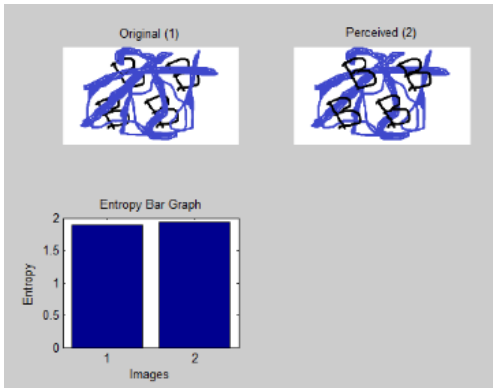
Amodal Completion

### 3.3 SURFACE FILLING-IN

David Marr’s 2&1/2D theory [11] suggests that in a normal situation, edges of the objects are detected first, and then, assuming continuity in brightness for the segmented object, our brain tries to fill in the whole surface to make a smooth texture. However, while the brightness of edges is not sharply different from the background and there is some fuzziness in the edges, surface filling-in occurs even though the real surface is absent. In these cases of illusory surface filling-in, our brain adds extra information to keep the continuity in surfaces for those edge-segmented objects so that they behave like real things in the world. Watercolor illusion and Neon color spreading [12] (From Figure 3 (Left) and (Right), respectively) can be categorized into this area, where surface filling-in is induced by the brain despite the retinal inputs not having the information.

**Figure 3:**
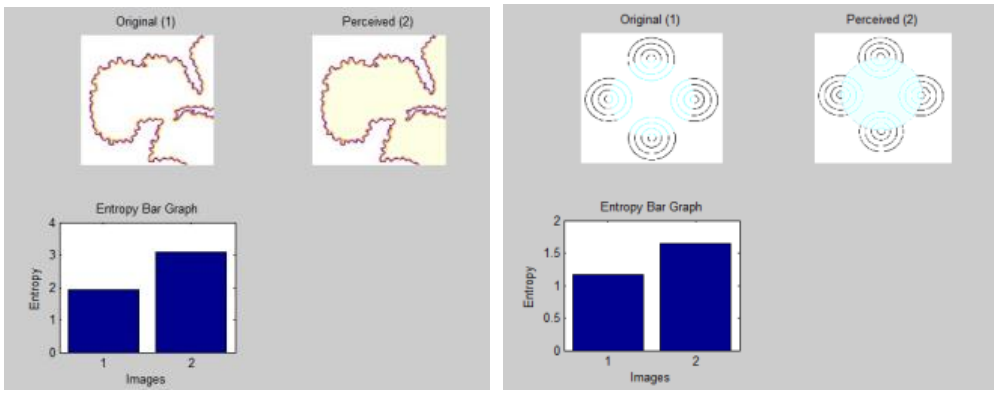
Watercolor illusion (Left) & Neon color spreading (Right)

### 3.4 BRIGHTNESS PERCEPTION

Extended Classical Receptive Field (ECRF) model of the retina has been implemented to describe the low-level brightness-contrast illusions, which outperformed the existing Difference of Gaussian (DoG) models [13]. In the ECRF model, there is one more extended disinhibitory surround in DoG. However, in reality, there are two types of brightness illusions, namely brightness contrast and brightness assimilation. Neither of them is well described or generalized by the ECRF models. From the perspective of the extra information insertion by the brain, the object with the same brightness differs perceptually when placed before backgrounds with different brightness levels. This implies that one extra brightness information has been added to the perceived images, increasing the overall entropy of the resultant images. Thus, the increase in entropy for illusory images is a generic phenomenon that can be ascribed to all optical illusions.

#### Simultaneous Brightness Contrast

The brightness of the backgrounds induces this effect. On brighter and darker backgrounds, the same object appears darker and brighter, respectively. Figure 4 (Left) illustrates this with an increase in entropy.

**Figure 4:**
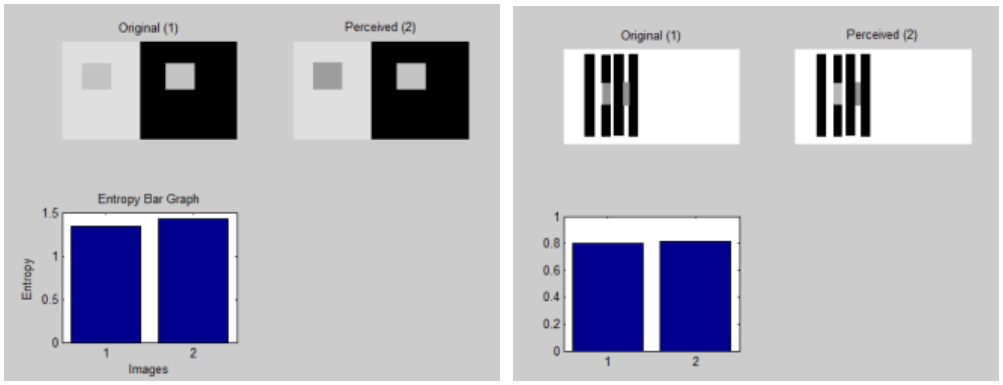
Simultaneous Brightness Contrast (Left) & Whiteness Illusion (Right)

#### White Illusion

In 1979, White [14] noticed that the brightness of objects surrounded by dark or bright borders does not follow the same logic as brightness contrast; rather it behaves just opposite to this convention. An identical grey bar looks brighter in the black strip than in the white strip. Figure 4 (Right) depicts this illusion where the entropy is higher in the illusory image than in the real input.

#### Checker Board Illusion

White’s explanation that the inhibition of the receptive fields along the spatial structure of the bar is stronger than across does not hold well for this Checkerboard illusion. This phenomenon can be termed brightness assimilation where the darker neighborhood perceptually diminishes the brightness of the object, so that in a lighter neighborhood the object astonishingly looks brighter. Figure 5 (Left) shows the original and perceived images along with the entropy measurements.

**Figure 5:**
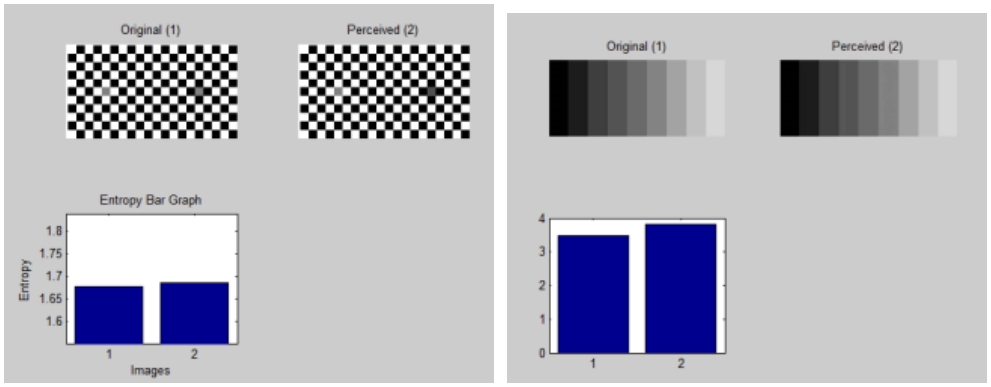
Checker Board Illusion (Left) & Mach Band Illusion (Right)

#### Mach Band Illusion

Almost a century before neurophysiological experiments began to evolve, Ernst Mach anticipated the center-surround receptive field structure [15]. He observed that when the brightness of a bar started to change from dark to light, humans observed a dark bar and when the brightness stopped changing, a lighter bar was observed. This extra dark and light bar information is added, increasing the entropy of the visual outputs, as seen in Figure 5 (Right).

### 3.5 SIZE PERCEPTION ILLUSION

Whenever we observe lines of the same length oriented vertically and horizontally, we perceive the vertical bar to be longer than the horizontal one. For this phenomenon, our brain inserts extra information into the vertical bar to increase the overall entropy of the perceived outcome (see Figure 6 (Left)).

**Figure 6:**
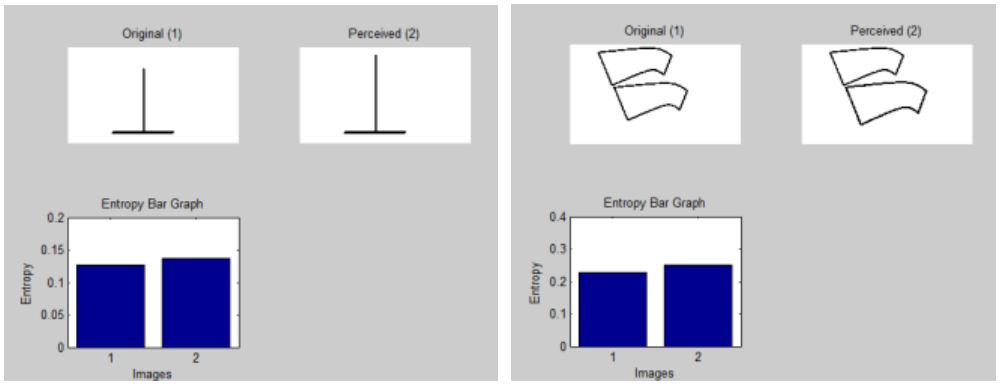
Vertical-Horizontal Illusion (Left) & Curved Bar Illusion (Right)

In Figure 6 (Right), the original retinal input has two curved bars of equal length, but we perceive the lower bar as larger than the upper one as described in the perceived image. In this case as well, the perceived image has higher entropy than the original one because some extra perimeter information is introduced in the lower bar.

### 3.5 HERMANN GRID ILLUSION

Ludimar Hermann in 1870 [16] discovered that some grey blobs appear between the black square neighborhoods, separated by the white regions, as shown in Figure 7. Similarly here, the perceived image exhibits greater entropy than the original, owing to the appearance of grey blobs at the intersections where the white lanes cross.

**Figure 7:**
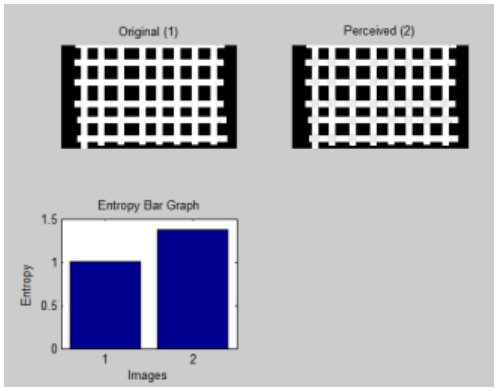
Hermann Grid Illusion

## CONCLUSION

All visual illusion explanation methods try to accommodate the logic of specific illusory patterns in a disintegrated way. However, if we consider our brain to be a workplace of synchronous neurons, there should be a uniformity and integrity with respect to all the illusion explanations. So, this increase in entropy is found to be a good generic explanation for all of the illusions. Though our brain tries to minimize energy consumption by keeping everything in low entropy, this may not always be possible because, to survive in a real environment, we should search for global feature similarity using local inputs to have a better perspective.

## Notes

### Competing Interest Statement

The authors have declared no competing interest.

